# Extracting the Research Goal from Biomedical Abstracts

**DOI:** 10.1101/2025.09.19.677266

**Authors:** Mariana Neves, Ines Schadock, Daniel Butzke, Gilbert Schönfelder, Bettina Bert

## Abstract

Extracting the research goal of a publication can potentially support researchers when searching the biomedical literature. Systems can make use of this information for various tasks, e.g., query processing or matching and ranking the candidates. However, this is a very subjective and complex task, which might involve various semantic types and may vary depending on the research area. Previous work have ventured in this area, but as far as we know, no specific dataset is yet available. We reused seven reviews from the European Commission with annotations about the research goal for more than 2.8k articles. We compiled the RG4C dataset, which we then used to fine tune a model for the automatic extraction of four criteria: “Field of Application”, “Disease Area”, “Disease Feature”, and “Biological Endpoint”. We obtained an overall f-score of 0.59, with results for each criterion ranging from 0.35 to 0.83. The RG4C dataset is available at: https://www.kaggle.com/datasets/marianaln/eu-qa-complete. Our source code is available at: https://www.kaggle.com/code/marianaln/eu-qa-features/notebook

## Introduction

For biomedical literature research, PubMed (https://pubmed.ncbi.nlm.nih.gov/) is the most prominent literature database and it currently contains more than 37 millions citations. When searching in PubMed, researchers usually rely on rather simple and short queries composed of keywords, MeSH terms and boolean operators (e.g., AND, OR) [1]. Previous work succeeded in representing queries in a more structured way with the aim to improve results. For instance, the PICO elements are widely used for evidence-based medicine [2].

The rampant growth in the number of biomedical publications generated a rise in the development of tools to support researchers for mining the literature [3, 4]. In spite of this, tailored tools are usually necessary for specific tasks, e.g., evidence-based medicine or genomics [3], which might include the precise extraction of the research goal. More recently, the development of large language models (LLMs) created new solutions for many natural language processing (NLP) tasks, such as question answering or summarization [5].

Accurately processing the research goal is essential when carrying out a precise search, e.g., for retrieving potentially relevant articles, or ranking and filtering the candidates [3]. A research goal can be represented in a rather structured, e.g., the PICO elements [2], or unstructured manner, e.g., as a sentence [6], a reference article [7] or a question [8].

Defining a research goal is a complex and subjective task that may involve a variety of dimensions and concepts [9]. In a previous study [10], we asked three experts to highlight the sentences in four biomedical abstracts that (to their opinion) contains the research goal. We show an example for one [11] of the articles in Table 1. The sentences with full agreement described the human LRRK2, its mutations (R1441C and G2019S), which might cause Parkinson’s disease (PD), and the potential of LRRK2 transgenic mice for a better understanding of PD. The sentences for which two experts agreed discussed the phenotype that may occur in mice with one of these mutations and cited the effect of these mutations. Finally, sentences selected by only one of the experts provided general background information about the mutation of the LRRK2 gene for PD, presented the state of the art on mice experiments for this particular research goal, as well as detailed results on the observations of the proposed experiments. From the eight sentences in this article, including the title, the three experts agreed on two sentences, and two of the experts agreed on two others. This illustrates the complexity of deciding which parts of the abstract are relevant for the research goal and the ones that could probably be ignored, since they only provide general background information on the topic or details about the outcome of the experiments.

**Table 1.**
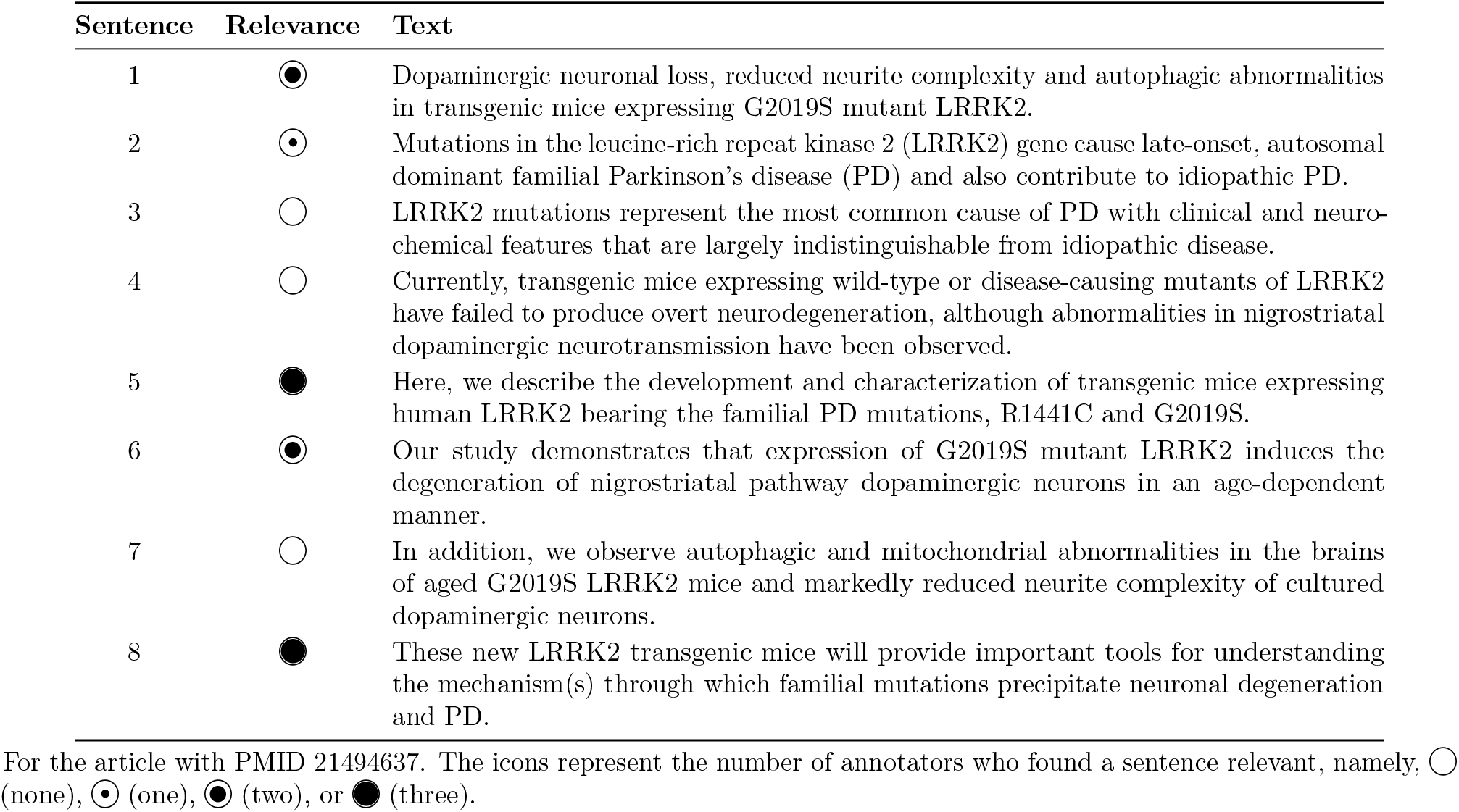
Relevance of an abstract’s sentences for the research goal, according to three annotators.

We are not aware of many previous efforts to extract the research goal in biomedical abstracts (cf. Related work). While some available corpora could be reused for this task, no comprehensive corpus seems to have been published for this aim.

In the present work we reused the recently published reviews from the European Commission (EC) for alternative methods to animal testing, i.e., mostly *in vitro* and *in silico* studies (cf. RG4C Dataset). Each review focuses on one topic (a disease) and includes a spreadsheet with information about the methods and research goal for the screened articles. Altogether, the seven reviews comprise a collection of more than 2.8k articles. We represented the research goal based on four criteria from the spreadsheets, namely “Field of Application”, “Disease Area”, “Disease Feature”, and “Biological Endpoint”. We compiled the RG4C dataset (Research Goal 4 Criteria) and trained a text generation model for their detection in abstracts. (We use the term “abstract” also when referring to the title and abstract of an article.)

We summarize below the main contributions of this article: (i) a dataset of more than 2.8k PubMed abstracts with annotations about the research goal (cf. RG4C Dataset), including an assessment of its quality (cf. Discussion); (ii) a fine-tuned model with the dataset (cf. Detection of the criteria); (iii) an evaluation of two LLMs and two tools for named-entity recognition (NER) (cf. Results). and (iv) a comprehensive analysis of the predictions of the fine-tuned model (cf. Discussion).

## Related work

### Corpora for the research goal

As far as we know, the only previous publication using the biomedical reviews was our preliminary experiments for the Field of Application criterion [12]. We modeled it as a text classification task, normalized similar labels, and grouped infrequent ones into a common label. The overall f-score was 0.57.

Previous work that tried to model the research goal in a structured way are still rare. One example is the EBM-NLP corpus, which contains 5k abstracts of clinical randomized controlled trials which were annotated with the PICO elements [13]. However, this schema is not suitable for articles not belonging to the clinical domain.

Some corpora contain potentially relevant annotations, but are limited to one field of research or one criterion. The LitCovid corpus for COVID-19 addressed the automatic classification of articles on pre-defined topics, e.g., “Case Report” or “Diagnosis” [14]. While some of them might overlap with the Field of Application criterion, it constitutes only one aspect of the research goal. A more recent corpus aimed at identifying four entity types related to cancer genomics [15], e.g., perturbing action, context, phenotype, and effect. The “perturbing action” and “phenotype” entities could overlap with the Biological Endpoint and Disease Feature criteria. However, the corpus is small (800 articles) and limited to cancer genomics. Further, the Hallmarks of Cancer (HoC) corpus originally contained 10 categories [16], which were later extended to a taxonomy of 37 classes [17]. However, most categories are specific for cancer research.

Two other corpora with similarities with the dataset that we compiled are the BioMRC [18] and the BioRead [19], both developed for machine reading comprehension (MRC). While they are certainly good benchmarks for the task, and some of the questions might refer to information from the research goal, the latter was not the aim of the corpora. Finally, our SMAFIRA-c dataset [9] comprises around 400 articles and contains eight categories for the stages of research, e.g., “target discovery” and “biological function”. Some of these categories overlap with the Field of Application criterion, however, the dataset is rather short for training purposes.

A variety of corpora were annotated to support NER, either for various semantic types, e.g. CRAFT [20], or for specific ones, e.g., gene/proteins [21], chemicals [22], or diseases [23]. While these are very comprehensive corpora with annotations for practically all mentions of the entities, it is uncertain which of these entities are actually part of the research goal. The latter problem was recently researched in [24], in which the authors addressed the distinction between focus and background entities, but limited to pathogens.

### Automatic detection of the research goal

Some of the criteria in a research goal overlap with available NER corpora (cf. above) and might be detected with the available NER tools for the biomedical domain. For instance, tools for extracting gene and diseases, e.g., BERN2 [25] or PubTator [26], could be used for the Biological Endpoint and Disease Area criteria, respectively. However, the Field of Application criterion could hardly be extracted with the available NER tools (cf. examples in Table 2), since the annotations do not seem to belong to a particular entity type. The Disease Feature and the Biological Endpoint criteria seem to be a mix of various entity types, such as anatomical parts (e.g., “aortic valve”), processes (e.g., “gene expression”), gene/protein names (e.g., “arid3b”), and even phrases (e.g., “least effective concentration and the half-life of the drug”). Further, not every entity type that is cited somewhere in the abstract is part of the research goal, since it depends in which context it was cited. In spite of the seemingly unsuitability of NER tools for our task, we still present an evaluation of two NER tools (cf. Results).

**Table 2.**
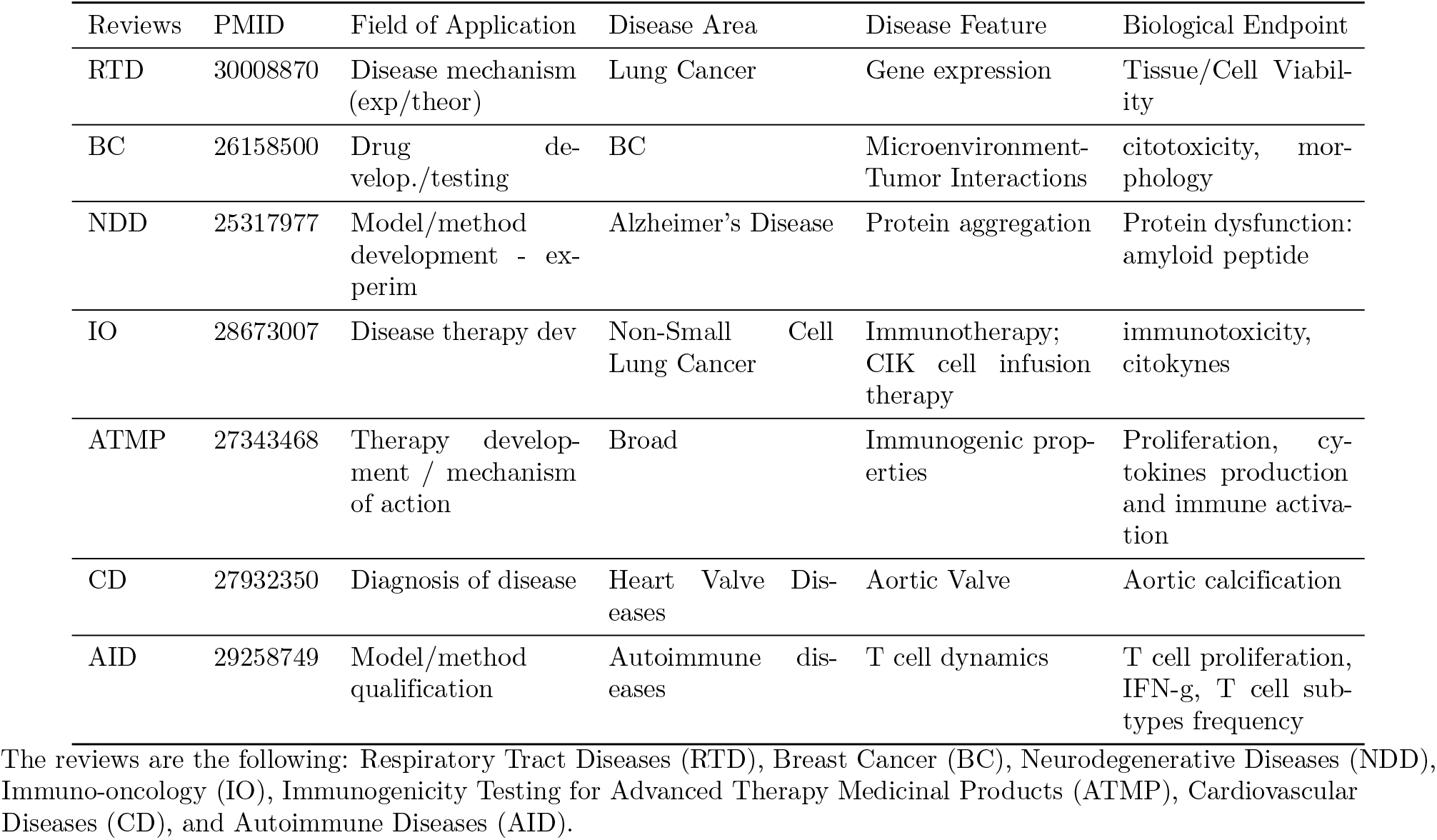
Examples of annotations for the four criteria in the seven reviews.

The current state-of-the-art methods for NLP heavily rely on LLMs [5]. Recent tools developed for NER, text classification (TC), and extractive question answering (QA) tasks usually rely on encoder-only LMs, e.g., AIONER [27] and LinkBERT [28]. However, this approach is not suitable for detecting the criteria, since the annotations do not include the corresponding text spans. Further, while the Field of Application criterion could be modeled as TC [29], the other criteria have far too many distinct annotations.

Given an appropriate prompt, various LLMs, e.g., Llama-3 (https://github.com/meta-llama/llama3) or GPT-4o (https://openai.com/index/gpt-4o-system-card/), are suitable for many NLP tasks, e.g., zero-shot or one-shot approaches [5]. However, results might not be comparable to those from tools which were fine-tuned for the particular task. We evaluated two LLMs for zero-, one-, and two-shot approaches and fine-tuned T5 [30] and BART [31] with the RG4C dataset (cf. Detection of the criteria).

## RG4C Dataset

In this section we introduce the biomedical reviews, describe how we processed the corresponding spreadsheets and compiled a dataset to train and evaluate our model. As opposed to some previous corpora (cf. Related work), the reviews contain document-level annotations and many of the annotations might not occur in the text of the article. It can be regarded as a structured summary of the articles and its size (around 2.8k) seems satisfactory for training purposes. Further, the four criteria provide a good representation of the research goal.

### Overview of the reviews

We reused the data of seven reviews from the the JRC’s EU Reference Laboratory which were published in the period of Sep/2020 to Nov/2022. These are reviews about alternatives to animal testing on the topic of “Advanced Non-animal Models in Biomedical Research”, namely, Respiratory Tract Diseases (RTD) [32], Breast Cancer (BC) [33], Neurodegenerative Diseases (NDD) [34], Immuno-oncology (IO) [35], Immunogenicity Testing for Advanced Therapy Medicinal Products (ATMP) [36], Cardiovascular Diseases (CD) [37], and Autoimmune Diseases (AID) [38]. Each review contains a corresponding spreadsheet with the curated data, e.g., experimental methods, diseases, and field of application. It also includes metadata for each article, e.g., title, DOI (Digital Identifier of an Object), authors, and the publication year. While most of the spreadsheets contain roughly the same columns, we noticed a few differences across them (cf. Supporting Information S1.1).

### Data model

We focused on four columns (i.e., criteria), namely, “Field of Application”, “Disease Area”, “Disease Feature”, “Biological Endpoint”. They all comply with the following: (a) occur in all seven reviews; (b) describe important facts from the research goal; (c) and have (mostly) free text in their annotations. However, they sometimes contain unspecific annotations, e.g., “broad” (for Disease Area), “no specific feature” or “na” (not applicable). We present some examples of the four criteria in Table 2. We provide a short definition of each criterion below, which was compiled from the various descriptions across the reviews:

- Field of Application: the main scientific aim of the article, or the application of the described model or method.
- Disease Area: area of the disease under study.
- Disease Feature: specific features of the particular disease which are under study.
- Biological Endpoint: characteristics of a model or changes on it for a particular disease feature.

### Data processing

From each of the reviews, we automatically extracted the four criteria above, along with the Digital Object Identifier (DOI). These are sometimes represented by different columns names in the respective spreadsheet (cf. Supporting Information S1.2). We obtained 3,048 DOIs, namely, 284 for RTD, 935 for BC, 567 for NDD, 542 for IO, 88 for ATMP, 449 for CD, and 183 for AID. We automatically queried PubMed with the DOIs to find the respective citations. For 67 DOIs, no citation could be found, which we manually checked and found a citation for 25 of them. After this manual post-processing, we ended up with 3,004 citations. After removing missing citations, wrongly retrieved PubMed identifiers (PMIDs), and duplicated PMIDs, we ended up with 2,853 PMIDs (cf. Supporting Information S1.3). For these articles, we retrieved the four criteria from the respective spreadsheets. To further improve their suitability for our experiment, we applied some procedures to the original annotations (cf. Supporting Information S1.4).

### Statistics of the dataset

We present an overview of the annotations in the dataset in Table 3. These values correspond to all distinct annotations for the corresponding criterion, even when referring to the same concept.^1^ Most of the annotations appear only once in the dataset, except for the ones of the FA criterion (cf. Supporting Information S1.5). We split the dataset into 10 folds in a structured way, i.e., such that each review is roughly equally represented in each fold. This was meant to obtain 10 folds of training/development (90%) and blind test (10%) sets. We divided the training/development sets further to create training (90%) and development (10%) sets. This is the data that we used in our experiments.

**Table 3.** Statistics of the RG4C dataset.

| Reviews | No. Articles | Field of Application | Disease Area | Disease Feature | Biological Endpoint |
| --- | --- | --- | --- | --- | --- |
| RTD | 270 | 19 | 11 | 37 | 43 |
| BC | 880 | 11 | 8 | 25 | 1,589 |
| NDD | 487 | 10 | 3 | 9 | 44 |
| IO | 518 | 11 | 120 | 43 | 907 |
| ATMP | 88 | 7 | 25 | 11 | 115 |
| CD | 436 | 3 | 9 | 15 | 289 |
| AID | 181 | 5 | 48 | 22 | 462 |
| Overall | 2,853 | 42 | 210 | 142 | 3,153 |

We present the number of articles (No. Articles column) and the number of distinct annotations for each criterion. Values are shown per review and for the complete dataset. The sum of one column might differ from the overall value due of duplicates across the reviews. The reviews are the following: Respiratory Tract Diseases (RTD), Breast Cancer (BC), Neurodegenerative Diseases (NDD), Immuno-oncology (IO), Immunogenicity Testing for Advanced Therapy Medicinal Products (ATMP), Cardiovascular Diseases (CD), and Autoimmune Diseases (AID).

## Detection of the criteria

In this section we describe our method for the automatic detection of the criteria, as well as the baselines that we considered.

### Fine-tuning with the RG4C dataset

We ran some experiments for automatically extracting the four criteria from an abstract. We relied on generative LMs, namely, T5 [30] and BART [31], which achieved good results for many downstream tasks. We fed the system with a suitable prompt for extracting the criteria, similar as carried out in [39].

We relied on a single prompt for all four criteria (cf. Table 4 and the algorithm in Supporting Information S2.1). The prompt contained the following elements: (a) a context, which consists of the title and abstract; (b) a question, which we manually created for each criterion; and (c) an answer for each question. This is processed by the tokenizer and fed to a Transformer model.

**Table 4.** Example of context (abstract text), questions and respective answers.

|  |  |
| --- | --- |
| Context | Lung Adenocarcinoma Cell Responses in a 3D in Vitro Tumor Angiogenesis Model Correlate with Metastatic Capacity<br>Many tools from the field of tissue engineering can be used to develop novel model systems to study cancer. [truncated] |
| Question | Answer the questions at the end based on the text.<br>1.What is the scientific aim of the work?<br>2.What is the specific disease type or pathological condition?<br>3.What is the most characteristic symptom/feature of the disease/pathology?<br>4.Which biological process was studied? |
| Answers | 1.Disease mechanism (exp/theor)<br>2.lung cancer<br>3.angiogenesis<br>4.Vascularisation |

Example for the article with PMID 33418731.

We tried ten distinct prompts (cf. Supporting Information S2.2) and multiple questions for each criterion (cf. Supporting Information S2.3), namely, three for Field of Application, two for Disease Area, four for Disease Feature, and four for Biological Endpoint. We used the following values for the hyperparameters: maximum length of 512, epochs of 5, 10 and 15, and batch size of 8. We ran all our experiments in the Kaggle platform (https://www.kaggle.com/) (GPU T4×2) and were limited by their memory constraints. For instance, we could only try T5-small (https://huggingface.co/google-t5/t5-small) and BART-Base (https://huggingface.co/facebook/bart-base). The model generates three answers for each question. We considered all of them for the Biological Endpoint criterion, which often contains multiple annotations, but only the first one for the other criteria.

### Few-shot evaluation

We tried two LLMs, namely, GPT-4o-mini using the API from OpenAI (https://openai.com/), and Llama (llama-3.3-70b-versatile) using the free API from GROQ (https://groq.com/). We queried the models with the best performing questions, which were identified with experiments during the fine-tuning approach.

The user message consisted of the following text: “Answer the following four questions below based on the text provided at the end.”, followed by the four questions, as presented in Table 4. Subsequently, we stated that the answers should be short (“The answers should be short, up to 100 characters.”), followed by the reference to the context (“This is the text:”), which we concatenated at the end. The response from the LLMs consisted of four itemized answers, which we processed accordingly. We provide an example of the complete prompt and the answers in Supporting Information S2.4. For few-shot evaluation, we randomly picked one or two examples from the training data.

We used a distinct prompt for the Field of Application criterion, since the results with the above prompt were very poor (results not shown). This criterion has fewer annotations and uses a rather limited number of terms (cf. Table 3). Therefore, we modeled it as a text classification task with the following prompt: “Assign one or more of the categories below based on the text provided at the end.”, followed by the list of categories and the context at the end. From the 42 annotations for the Field of Application criterion, we normalized some duplicates, e.g., “Diagnosis of disease” and “Diagnosis of Disease”, and ended up with 28 categories. The format for the response varied in many cases, and we processed them accordingly. We provide an example of the complete prompt and the responses in Supporting Information 2.4.

### Evaluation of NER tools

NER tools could potentially be used for detecting three of the criteria (cf. Section Related work). We tested this hypothesis by evaluating two NER tools, namely, BERN2 [25] and PubTator [26], and tagged all articles in the dataset with the two tools. Subsequently, we analyzed the overlap between the entity types of the predictions with the annotations for each criterion to identify which entity types best correlate with each criterion (cf. Supporting Information 2.5). Based on this analysis, we defined a mapping for each of the NER tools, i.e., the respective disease type mapped to the Disease Area and Disease Feature criteria and the respective gene/protein type for the Biological Endpoint criterion (cf. Supporting Information S2.5)

We ranked the predictions from the tools from the most to the least frequent one. Our hypothesis was that predictions that occur more frequently are more likely to be part of the research goal. Finally, we considered just the most frequent one for Disease Area and Disease Feature, and the top three ones for Biological Endpoint.

## Results

We show results in terms of precision, recall, and f-score. For each criterion, we compared the annotations from the dataset with the answers returned by the systems (i.e., predictions). Since we are comparing free-text answers, we tried three string similarity methods for the comparison: (i) strict, i.e., perfect match between the dataset annotations and the prediction; (ii) overlap, i.e., whether the dataset annotation is included in the prediction, or vice-versa; (iii) and Levenshtein distance [40] between the dataset annotation and the prediction.

In Table 5 we present the results for the fine-tuned model and the LLMs. The best results with fine-tuning were obtained with BART-Base, 10 epochs and a batch size of 8. For the Levenshtein distance, we adopted a threshold of 0.6 after a manual analysis of the predictions (cf. Section Error analysis and threshold for the Levenshtein distance). The best performing prompt and questions are the ones presented in Table 4.

**Table 5.**
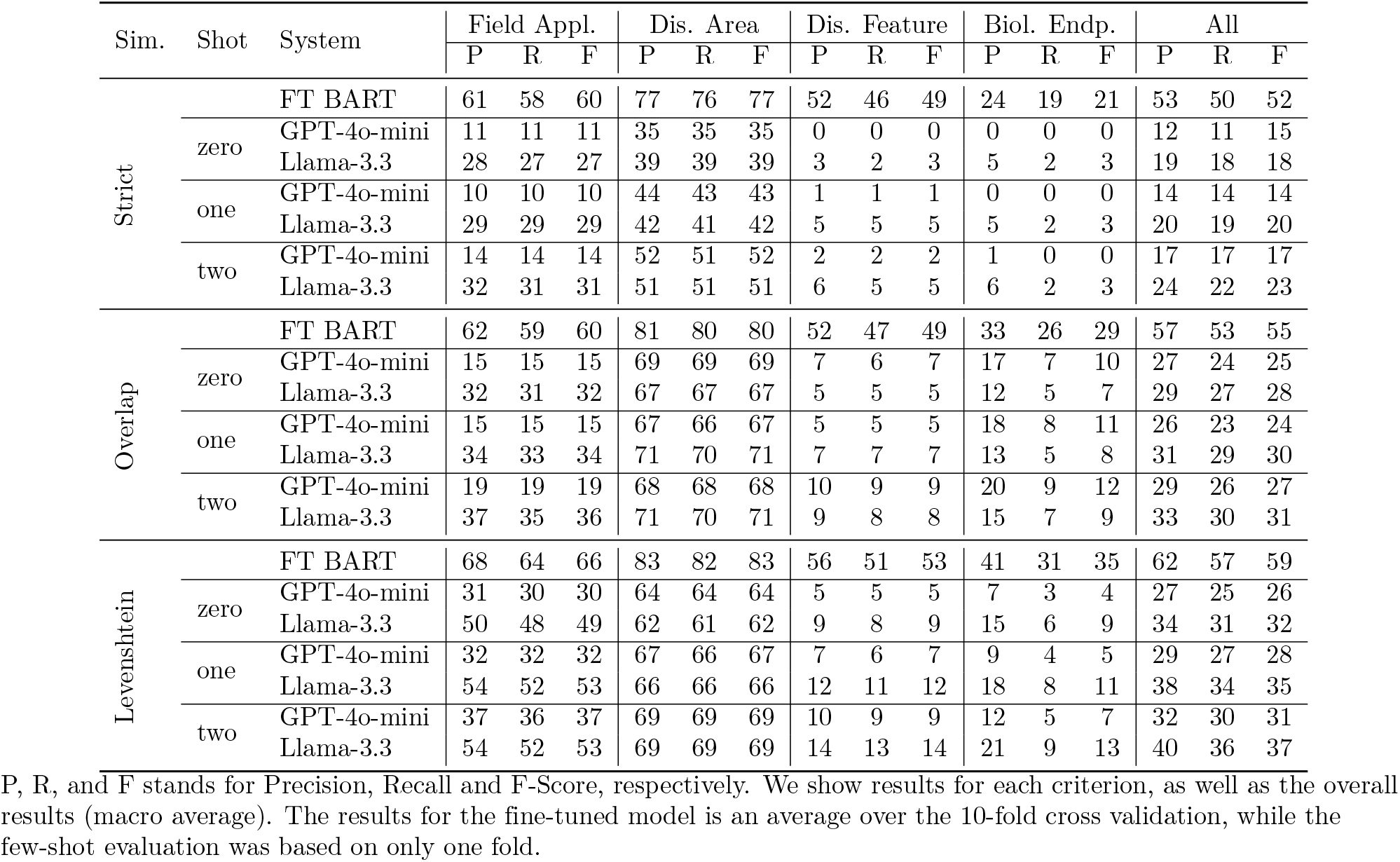
Best results for the fine-tuned (FT) model with BART, and few-shot with GPT-4o-mini and Llama-3.3.

In general, results were higher for Disease Area and lower for Biological Endpoint. The fine-tuned model outperformed the LLMs for all criteria, metrics, and similarity degrees. The performance of GPT-4o-mini was usually lower than results from Llama3.3, with some few exceptions. One reason might be that GPT-4o-mini returned rather long answers, and the additional characters were strongly penalized during the evaluation, specially by the Levenshtein distance. We discuss the errors made by the fine-tuned model in Section Error analysis and threshold for the Levenshtein distance.

We ran some ablation experiments for the fine-tuned model. The results were considerably better when using 10 epochs (cf. Supporting Information S3.1). We tried different questions for the criteria, but could not observe a consistent behavior (cf. Supporting Information S3.2), and since we used a multi-question prompt, one question might have been influenced by another question. We ended up adopting the questions presented in Table 4. We tried two language models, i.e., T5-Small and BART-Base, and the results for the latter were much better for all four criteria (cf. Supporting Information S3.3). Finally, we experimented with various prompts, but there was little difference in the results (cf. Supporting Information S3.4), and decided for the one presented in Table 4.

We computed f-scores for each criterion and review (cf. Supporting Information S3.5). There are some differences across the reviews, i.e., some criteria are harder to be predicted in some reviews, but we did not observe that one review outperformed the others for all four criteria. For instance, the scores for Biological Endpoint are rather low for Autoimmune Diseases and Respiratory Tract Diseases. Further, we could not observe a correlation with the number of distinct annotations per review and criterion (cf. Table 3). For instance, for the Biological Endpoint criterion, the Breast Cancer review has much more distinct annotations (around 1.5k) than Respiratory Tract Diseases (only 43). Nevertheless, the score for Breast Cancer is higher than for Respiratory Tract Diseases for this criterion.

Regarding the performance of the NER tools, there was not much difference between the two tools and results were rather poor (cf. Supporting Information S3.6), i.e., confirming that they are unsuitable for the task. However, both tools performed slightly better than GPT-4o-mini for the Disease Feature and Biological Endpoint criteria.

## Discussion

### Analysis of the dataset

We compiled a dataset using the original annotations from the spreadsheets of the seven reviews with a minimum of pre-processing (cf. Section RG4C Dataset). Since the reviews were not created for supervised learning purposes, the derived dataset does not comply with the quality standards usually carried out for corpora, e.g., annotation guidelines or inter-annotator agreement.

Even though the authors provided a short definition for each criterion, there is no detailed description on how the annotation was carried out, whether there was an agreement between the annotators, or whether a particular article was annotated by more than one annotator. Further, we assume that the full text of the articles was often taken into consideration. The annotations were often provided as free text, sometimes as abbreviations or acronyms, without a mapping to a particular ontology or database, for instance, for the disease names.

Despite the above observations, we believe that the reviews (and their corresponding spreadsheets) are a valuable resource due to various reasons: (a) its size (around 2.8k), which suitable for training purposes; (b) the variety of the annotations, since many can only be found in some articles or one review; (c) the comprehensiveness of the annotations, since all articles have annotations for the four criteria; and (d) the lack of a similar dataset with annotations for the research goal.

We observed many differences across the reviews. As presented in Table 3, some of them clearly contain much more articles (e.g., 880 for Breast Cancer) than others (e.g., 88 for Immunogenicity Testing for Advanced Therapy Medicinal Products). The number of distinct annotations per criterion varied considerably across the reviews, especially for the Disease Area and Biological Endpoint criteria. This shows the heterogeneity across the reviews, e.g., the differences in their annotation.

For each criterion, we analyzed the number of annotations that occur in more than one review (cf. Supporting Information S4.1). Around 70% of Field of Application can only be found in one review, mainly due to different writing of the same concepts. Additionally, some reviews have specific annotations for this criterion, e.g., Respiratory Tract Diseases. As expected, around 94% of the DA criterion occur in only one review, but some of them appear in more than one, usually the cancer-related ones, but also some others, e.g., “inflammatory bowel disease” and “psoriasis”. The Disease Feature and Biological Endpoint criteria also had around 90% of their annotations occurring in only one review.

### Error analysis and threshold for the Levenshtein distance

We carried out an analysis of the predictions for one of the test sets (from the 10 folds). This aimed not only to provide insights on the errors made by the fine-tuned model, but also to define the threshold for the Levenshtein distance. We validated all the pairs (i.e., dataset annotation versus prediction) based on a three-value scale: correct, partially correct, or incorrect. The definition of each of these values slightly varied for each criterion (cf. Supporting Information S4.2). We summarize the results of the manual evaluation in Table 6.

**Table 6.** Results of the manual evaluation.

| Criteria | correct | partial | incorrect | total |
| --- | --- | --- | --- | --- |
| Field of Application | 29 | 19 | 36 | 84 |
| Disease Area | 47 | 26 | 14 | 87 |
| Disease Feature | 33 | 33 | 60 | 126 |
| Biological Endpoint | 82 | 117 | 335 | 534 |

For all distinct pairs of dataset annotation and prediction for one test set (fold # 0) of the 10-fold cross-validation.

For the Field of Application criterion, all predictions were valid annotations for this criterion and included different versions of the same answer, e.g., “Drug develop/testing” and “Drug developm/testing”. For the Disease Area criterion, the overall impression of the predictions was that all of them do seem to be diseases. Some few exceptions were predictions such as “broad” and “general”, which are rather vague. Most of the predictions for Disease Feature seemed to belong to biomedical terms and could potentially be correct. However, similar to the ones for DA, some of them were vague, namely “na”, “None”, “exploratory/no specific feature”.

Finally, we observed a higher rate of incorrect matches for Biological Endpoint, probably because we allowed up to three answers for this criterion. The great majority of these predictions seemed to be valid biological terms, except for some occurrences of “Other”. Further, this was the only criterion for which we observed predictions that clearly belonged to another criterion, e.g., “Disease mechanism” and “Disease mechanism (exp/theor)” belong to Field of Application. There were many gene and protein names for this criterion, but given the the high number of pairs, we could not check all of them in details. When in doubt, we opted for labeling a pair as “incorrect”.

We calculated the Levenshtein distance for each pair of dataset annotation and prediction. The higher the threshold we defined for the maximum Levenshtein distance, the higher was the risk of including incorrect and partially correct matches. In our automatic evaluation (cf. Section) we considered a threshold of 0.6. This means that the true positives could potentially include 10%-40% of partially correct matches and 5%-10% of incorrect matches. We present the detailed results (precision, recall, and f-score) and the proportion of correct, partially incorrect and incorrect matches for each threshold as supplementary files. Based on our manual validation and the distance scores, we plotted in Figure 1 the proportion of each degree, i.e., correct, partially correct, or incorrect, for distances ranging from 0.0 (only perfect matches) to 1.0 (any difference allowed).

**Fig 1.**
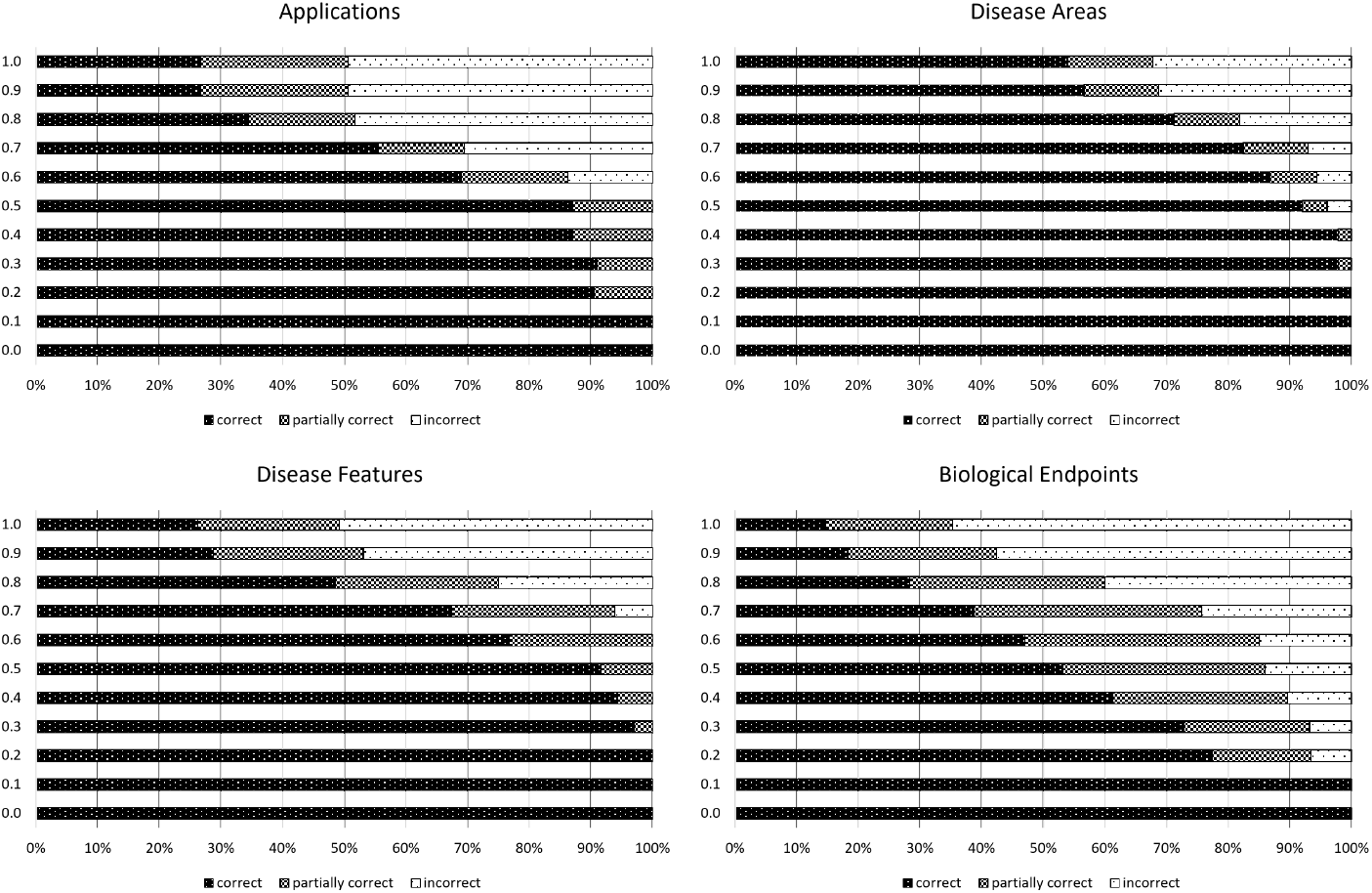
Rate of correct, partially correct, and incorrect answers (x-axis) that would be considered as true positives depending on the threshold for the Levenshtein distance (y-axis).

## Conclusion

We reused the data released in the reviews from the European Commission to create the RG4C dataset, which aims to represent the research goal of biomedical abstracts. As far as we know, this is the first attempt to model the research goal, since previous corpora were either limited to one topic or to one criterion. Even though the dataset is not comparable to available gold standard corpora, e.g., for the PICO elements, we believe that this is a first step to motivate more work in this area, and future efforts to improve our methods and the dataset. We hope that this dataset can be used as a benchmark for this task.

The fine-tuned model obtained reasonable results, given the difficulty of the task, i.e., the dataset on which it was trained contained document-level annotations. However, it outperformed the LLMs for all four criteria, as well as the two NER tools, i.e., BERN2 and PubTator. Future research could explore whether the results could have been further improved by fine-tuning other LLMs, e.g., Llama3, or even domain-specific LLMs.

## Supporting information

### Supplementary material

PDF file with various additional information.

## Author contributions

- Formal Analysis: Mariana Neves
- Funding Acquisition: Gilbert Schönfelder
- Methodology: Mariana Neves
- Project Administration: Bettina Bert
- Resources: Mariana Neves
- Software: Mariana Neves
- Supervision: Bettina Bert
- Validation: Mariana Neves, Daniel Butzke
- Visualization: Mariana Neves
- Writing, Review & Editing: Mariana Neves, Ines Shadock, Daniel Butzke, Bettina Bert, Gilbert Schönfelder

## Footnotes

1 For instance, “Model/method dev - exp” and “Model development - experim” were considered as distinct annotations, since we did no try to normalize them to a common one.

## References

1. Islamaj Dogan R, Murray GC, Névéol A, Lu Z. Understanding PubMed® user search behavior through log analysis. Database. 2009;2009:bap018. doi:10.1093/database/bap018.

2. Richardson S, Wilson M, Nishikawa J, Hayward R. The well-built clinical question: a key to evidence-based decisions. ACP journal club. 1995;123(3):A12–13.

3. Jin Q, Leaman R, Lu Z. PubMed and beyond: biomedical literature search in the age of artificial intelligence. eBioMedicine. 2024;100. doi:10.1016/j.ebiom.2024.104988.

4. Lu Z. PubMed and beyond: a survey of web tools for searching biomedical literature. Database. 2011;2011:baq036. doi:10.1093/database/baq036.

5. Tian S, Jin Q, Yeganova L, Lai PT, Zhu Q, Chen X, et al. Opportunities and challenges for ChatGPT and large language models in biomedicine and health. Briefings in Bioinformatics. 2024;25(1):bbad493. doi:10.1093/bib/bbad493.

6. Allot A, Chen Q, Kim S, Vera Alvarez R, Comeau DC, Wilbur WJ, et al. LitSense: making sense of biomedical literature at sentence level. Nucleic Acids Research. 2019;47(W1):W594–W599. doi:10.1093/nar/gkz289.

7. Mysore S, O’Gorman T, McCallum A, Zamani H. CSFCube - A Test Collection of Computer Science Research Articles for Faceted Query by Example. In: Vanschoren J, Yeung SK, editors. NeurIPS Datasets and Benchmarks; 2021. Available from: http://dblp.uni-trier.de/db/conf/nips/neurips2021db.html#MysoreOMZ21.

8. Fontelo P, Liu F, Ackerman M. ask MEDLINE: a free-text, natural language query tool for MEDLINE/PubMed. BMC Medical Informatics and Decision Making. 2005;5(1):5. doi:10.1186/1472-6947-5-5.

9. Butzke D, Dulisch N, Dunst S, Steinfath M, Neves M, Mathiak B, et al. SMAFIRA-c: A benchmark text corpus for evaluation of approaches to relevance ranking and knowledge discovery in the biomedical domain; 2020. Available from: 10.21203/rs.3.rs-16454/v1.

10. Neves M, Schadock I, Eusemann B, Schnfelder G, Bert B, Butzke D. Is the ranking of PubMed similar articles good enough? An evaluation of text similarity methods for three datasets. In: Demner-fushman D, Ananiadou S, Cohen K, editors. The 22nd Workshop on Biomedical Natural Language Processing and BioNLP Shared Tasks. Toronto, Canada: Association for Computational Linguistics; 2023. p. 133–144. Available from: https://aclanthology.org/2023.bionlp-1.11.

11. Ramonet D, Daher JPL, Lin BM, Stafa K, Kim J, Banerjee R, et al. Dopaminergic Neuronal Loss, Reduced Neurite Complexity and Autophagic Abnormalities in Transgenic Mice Expressing G2019S Mutant LRRK2. PLOS ONE. 2011;6(4):1–15. doi:10.1371/journal.pone.0018568.

12. Neves M, Butzke D, Grune B. Evaluation of Scientific Elements for Text Similarity in Biomedical Publications. In: Stein B, Wachsmuth H, editors. Proceedings of the 6th Workshop on Argument Mining. Florence, Italy: Association for Computational Linguistics; 2019. p. 124–135. Available from: https://aclanthology.org/W19-4515.

13. Nye B, Li JJ, Patel R, Yang Y, Marshall I, Nenkova A, et al. A Corpus with Multi-Level Annotations of Patients, Interventions and Outcomes to Support Language Processing for Medical Literature. In: Gurevych I, Miyao Y, editors. Proceedings of the 56th Annual Meeting of the Association for Computational Linguistics (Volume 1: Long Papers). Melbourne, Australia: Association for Computational Linguistics; 2018. p. 197–207. Available from: https://aclanthology.org/P18-1019.

14. Chen Q, Allot A, Leaman R, Islamaj R, Du J, Fang L, et al. Multi-label classification for biomedical literature: an overview of the BioCreative VII LitCovid Track for COVID-19 literature topic annotations. Database. 2022;2022:baac069. doi:10.1093/database/baac069.

15. Collins C, Baker S, Brown J, Zheng H, Chan A, Stenius U, et al. Text mining for contexts and relationships in cancer genomics literature. Bioinformatics. 2024;40(1):btae021. doi:10.1093/bioinformatics/btae021.

16. Baker S, Silins I, Guo Y, Ali I, Högberg J, Stenius U, et al. Automatic semantic classification of scientific literature according to the hallmarks of cancer. Bioinformatics. 2015;32(3):432–440. doi:10.1093/bioinformatics/btv585.

17. Baker S, Ali I, Silins I, Pyysalo S, Guo Y, Högberg J, et al. Cancer Hallmarks Analytics Tool (CHAT): a text mining approach to organize and evaluate scientific literature on cancer. Bioinformatics. 2017;33(24):3973–3981. doi:10.1093/bioinformatics/btx454.

18. Pappas D, Stavropoulos P, Androutsopoulos I, McDonald R. BioMRC: A Dataset for Biomedical Machine Reading Comprehension. In: Demner-Fushman D, Cohen KB, Ananiadou S, Tsujii J, editors. Proceedings of the 19th SIGBioMed Workshop on Biomedical Language Processing. Online: Association for Computational Linguistics; 2020. p. 140–149. Available from: https://aclanthology.org/2020.bionlp-1.15.

19. Pappas D, Androutsopoulos I, Papageorgiou H. BioRead: A New Dataset for Biomedical Reading Comprehension. In: Calzolari N, Choukri K, Cieri C, Declerck T, Goggi S, Hasida K, et al., editors. Proceedings of the Eleventh International Conference on Language Resources and Evaluation (LREC 2018). Miyazaki, Japan: European Language Resources Association (ELRA); 2018. Available from: https://aclanthology.org/L18-1439.

20. Bada M, Eckert M, Evans D, Garcia K, Shipley K, Sitnikov D, et al. Concept annotation in the CRAFT corpus. BMC Bioinformatics. 2012;13(1):161. doi:10.1186/1471-2105-13-161.

21. Smith L, Tanabe LK, Ando RJn, Kuo CJ, Chung IF, Hsu CN, et al. Overview of BioCreative II gene mention recognition. Genome Biology. 2008;9(2):S2. doi:10.1186/gb-2008-9-s2-s2.

22. Li J, Sun Y, Johnson RJ, Sciaky D, Wei CH, Leaman R, et al. BioCreative V CDR task corpus: a resource for chemical disease relation extraction. Database. 2016;2016:baw068. doi:10.1093/database/baw068.

23. Doğan RI, Leaman R, Lu Z. NCBI disease corpus: A resource for disease name recognition and concept normalization. Journal of Biomedical Informatics. 2014;47:1–10. doi:10.1016/j.jbi.2013.12.006.

24. Jimeno Yepes A, Verspoor K. Distinguishing between focus and background entities in biomedical corpora using discourse structure and transformers. In: Lavelli A, Holderness E, Jimeno Yepes A, Minard AL, Pustejovsky J, Rinaldi F, editors. Proceedings of the 13th International Workshop on Health Text Mining and Information Analysis (LOUHI). Abu Dhabi, United Arab Emirates (Hybrid): Association for Computational Linguistics; 2022. p. 35–40. Available from: https://aclanthology.org/2022.louhi-1.4/.

25. Sung M, Jeong M, Choi Y, Kim D, Lee J, Kang J. BERN2: an advanced neural biomedical named entity recognition and normalization tool. Bioinformatics. 2022;38(20):4837–4839. doi:10.1093/bioinformatics/btac598.

26. Wei CH, Allot A, Leaman R, Lu Z. PubTator central: automated concept annotation for biomedical full text articles. Nucleic Acids Research. 2019;47(W1):W587–W593. doi:10.1093/nar/gkz389.

27. Luo L, Wei CH, Lai PT, Leaman R, Chen Q, Lu Z. AIONER: all-in-one scheme-based biomedical named entity recognition using deep learning. Bioinformatics. 2023;39(5):btad310. doi:10.1093/bioinformatics/btad310.

28. Yasunaga M, Leskovec J, Liang P. LinkBERT: Pretraining Language Models with Document Links. In: Muresan S, Nakov P, Villavicencio A, editors. Proceedings of the 60th Annual Meeting of the Association for Computational Linguistics (Volume 1: Long Papers). Dublin, Ireland: Association for Computational Linguistics; 2022. p. 8003–8016. Available from: https://aclanthology.org/2022.acl-long.551.

29. Neves M. Detection of fields of applications in biomedical abstracts with the support of argumentation elements; 2024.

30. Raffel C, Shazeer N, Roberts A, Lee K, Narang S, Matena M, et al. Exploring the Limits of Transfer Learning with a Unified Text-to-Text Transformer. Journal of Machine Learning Research. 2020;21(140):1–67.

31. Lewis M, Liu Y, Goyal N, Ghazvininejad M, Mohamed A, Levy O, et al. BART: Denoising Sequence-to-Sequence Pre-training for Natural Language Generation, Translation, and Comprehension. In: Jurafsky D, Chai J, Schluter N, Tetreault J, editors. Proceedings of the 58th Annual Meeting of the Association for Computational Linguistics. Online: Association for Computational Linguistics; 2020. p. 7871–7880. Available from: https://aclanthology.org/2020.acl-main.703.

32. Commission E, Centre JR, Adcock I, Novotny T, Nic M, Dibusz K, et al. Advanced non-animal models in biomedical research : respiratory tract diseases. Publications Office of the European Union; 2020.

33. Commission E, Centre JR, Rossi F, Caforio M, Nic M, Dibusz K, et al. Advanced non-animal models in biomedical research : breast cancer. Publications Office of the European Union; 2020.

34. Commission E, Centre JR, Witters H, Verstraelen S, Aerts L, Miccoli B, et al. Advanced non-animal models in biomedical research : neurodegenerative diseases. Publications Office of the European Union; 2021.

35. Commission E, Centre JR, Romania P, Folgiero V, Nic M, Dibusz K, et al. Advanced non-animal models in biomedical research : immuno-oncology. Publications Office of the European Union; 2021.

36. Commission E, Centre JR, Canals J, Romania P P BM, Nic M, et al. Advanced Non-animal Models in Biomedical Research - Immunogenicity testing for advanced therapy medicinal products. Publications Office of the European Union; 2022.

37. Commission E, Centre JR, Capellini K, Fanni B, Gasparotti E, Vignali E, et al. Advanced non-animal models in biomedical research : cardiovascular diseases. Publications Office of the European Union; 2022.

38. Commission E, Centre JR, Otero M, Canals J, Belio-Mairal P, Nic M, et al. Advanced non-animal models in biomedical research : autoimmune diseases. Publications Office of the European Union; 2022.

39. Arannil V, Deb T, Roy A. ADEQA: A Question Answer based approach for joint ADE-Suspect Extraction using Sequence-To-Sequence Transformers. In: Demner-fushman D, Ananiadou S, Cohen K, editors. The 22nd Workshop on Biomedical Natural Language Processing and BioNLP Shared Tasks. Toronto, Canada: Association for Computational Linguistics; 2023. p. 206–214. Available from: https://aclanthology.org/2023.bionlp-1.17.

40. Navarro G. A guided tour to approximate string matching. ACM Comput Surv. 2001;33(1):31–88. doi:10.1145/375360.375365.

